# Aboveground impacts of a belowground invader: how invasive earthworms alter aboveground arthropod communities in a northern North American forest

**DOI:** 10.1101/2021.12.02.470883

**Authors:** Malte Jochum, Lise Thouvenot, Olga Ferlian, Romy Zeiss, Bernhard Klarner, Ulrich Pruschitzki, Edward A. Johnson, Nico Eisenhauer

## Abstract

Declining arthropod communities have recently gained a lot of attention with climate and land-use change among the most-frequently discussed drivers. Here, we focus on a seemingly underrepresented driver of arthropod-community decline: biological invasions. For ∼12,000 years, earthworms have been absent from wide parts of northern North America, but they have been re-introduced with dramatic consequences. Most studies investigating earthworm-invasion impacts focus on the belowground world, resulting in limited knowledge on aboveground-community changes. We present observational data on earthworm, plant, and aboveground-arthropod communities in 60 plots, distributed across areas with increasing invasion status (low, medium, high) in a Canadian forest. We analyzed how earthworm-invasion status and biomass impact aboveground arthropod community abundance, biomass, and species richness, and how earthworm impacts cascade across trophic levels. We sampled ∼13,000 arthropods, dominated by Hemiptera, Diptera, Araneae, Thysanoptera, and Hymenoptera. Total arthropod abundance, biomass, and species richness declined significantly from areas of low to those with high invasion status with reductions of 61, 27, and 18%, respectively. Structural Equation Models unraveled that earthworms directly and indirectly impact arthropods across trophic levels. We show that earthworm invasion can alter aboveground multitrophic arthropod communities and suggest that belowground invasions can be important drivers of aboveground-arthropod decline.

## Introduction

Recent reports on arthropod species richness, abundance, and biomass declines (Habel et al. 2016, Hallmann et al. 2017, Seibold et al. 2019) have triggered concern about “the little things that run our world” (Wilson 1987) and the consequences of their loss. Even though the situation might not be equally bad for all taxa and ecosystem types (van Klink et al. 2020), the sheer number and extent of the reported negative trends, together with the fact that we simply do not have enough long-term datasets to establish such trends across all taxa and ecosystems (Stuart et al. 2010, Eisenhauer et al. 2019a, Eisenhauer and Hines 2021), is worrying. With arthropods contributing to central ecosystem processes and services (Noriega et al. 2018), their loss will have unprecedented consequences for ecosystems and human societies.

In order to halt or reverse arthropod decline, we need to understand the underlying drivers. Given their importance as broad global-change drivers (Díaz et al. 2019), it is unsurprising that climate and land-use change are prominent examples (Deutsch et al. 2008, Habel et al. 2016, Lister and Garcia 2018, van Klink et al. 2020). However, though underrepresented in research on arthropod declines, other drivers might still play an important role depending on the focal ecosystem and community. Here, we focus on one potentially underappreciated driver of arthropod decline: the invasion of a belowground ecosystem engineer, earthworms (Jones et al. 1994).

Although commonly perceived as having mostly positive impacts on their environment (Blouin et al. 2013, Van Groenigen et al. 2014), earthworms can transform invaded ecosystems (Hendrix and Bohlen 2002) that are not able to deal with their impacts on the ecosystem’s physical, chemical, and biological properties (Bohlen et al. 2004, Ferlian et al. 2020, Jochum et al. 2021). Earthworm invasion is a global problem (Hendrix et al. 2008). One region with both, particularly severe impacts and a lot of research on the consequences, is northern North America. Here, most earthworm species present today have been absent since the last glaciation and have only been re-introduced a few hundred years ago (Bohlen et al. 2004, James and Hendrix 2004).

Earthworm invasion alters soil abiotic conditions (Bohlen et al. 2004, Ferlian et al. 2020), plant communities (Nuzzo et al. 2009, Craven et al. 2017, Fleri et al. 2021), and soil fauna (Shao et al. 2017, Ferlian et al. 2018, McCay and Scull 2019, Jochum et al. 2021). Moreover, there are reports of consequences for aboveground vertebrates, such as salamanders, birds, and deer (Eisenhauer et al. 2019b, Frelich et al. 2019). There also are some aboveground-invertebrate studies, but these mostly focus on litter-dwelling fauna (Burtis et al. 2014, McCay and Scull 2019). With invasive earthworms impacting soil abiotic conditions, soil fauna, plants, and litter-dwelling arthropods, the open question is whether and how their invasion impacts aboveground, vegetation-dwelling arthropods, and if these changes cascade across trophic levels. For example, earthworms could directly serve aboveground arthropods as a food resource (King et al. 2010), or indirectly affect them via altered habitat structure, resource availability (leaf litter), or plant communities (Suárez et al. 2006, Nuzzo et al. 2009). We used earthworm, plant, and aboveground-arthropod community data from a Canadian forest to investigate i) whether belowground invasion by earthworms changes aboveground-arthropod communities and, using Structural Equation Models (SEMs), ii) how earthworms directly and indirectly impact higher trophic levels mediated by plants, herbivores, and detritivores. We expected invasive earthworms to decrease the abundance, biomass, and diversity of aboveground-arthropod communities via cascading effects across trophic levels (Scherber et al. 2010, Frelich et al. 2019).

## Material and Methods

We studied a south-facing forest slope above the Northwestern shore of Barrier Lake, Kananaskis Valley, Alberta, Canada (51°02’6’’N, 115°03’54’’W, ∼1450 m a.s.l). The forest is dominated by trembling aspen (*Populus tremuloides)* interspersed with balsam poplar (*Populus balsamifera)*, with a dense understorey vegetation and a grey luvisol soil. It has a long history of earthworm-invasion research (Scheu and Parkinson 1994, Eisenhauer et al. 2007, Jochum et al. 2021). We combine community data on earthworms, plants, and aboveground arthropods sampled in June and July 2019 on observational plots of the “EcoWorm” project (described in Eisenhauer et al. (2019)). After verifying earthworm-invasion status along the slope, we established 20 1×2m plots in each of three invasion-status areas: low, mid, high invasion (n=60 plots, **Appendix Fig. S1**). These categories differed significantly in earthworm abundance, biomass, species richness, and functional-group richness (**Appendix Fig. S2 and S3**). Thus, we focused on invasion status as the main predictor and show responses to earthworm biomass in the appendix. We used 1 m^2^ for plant-community assessments and the other half plot for arthropod (1 m^2^) and earthworm sampling (0.25 m^2^, **Appendix Fig. S4**). We identified every plant species and estimated total plant cover using a modified decimal scale (Londo 1976 ; details in **Appendix section 1**).

Earthworms were extracted using a combination of hand sorting and mustard extraction. All individuals were identified to species level, assigned to a functional group, and their fresh mass was assessed (**Appendix section 1**). We sampled aboveground arthropods using a vacuum suction sampler. All collected animals were hand-sorted, identified to (morpho-)species, assigned to a trophic feeding guild (see **Appendix section 2** for details, **Figures S5 and S6, and Table S1**), and their fresh biomass was estimated (**Appendix section 3**; Mercer et al. 2001, Wardhaugh 2013, Sohlström et al. 2018). We calculated abundance, biomass, and species richness of the total arthropod community, and, separately, for herbivores, omnivores (combining all mixed-diet feeding guilds), predators, detritivores, and parasitoids. While abundance and biomass were calculated based on all individuals (excluding mites and springtails), species richness was calculated based on adults only.

Data analysis was done in R version 3.6.3 (R Core Team 2020). We assessed arthropod community responses to invasion using earthworm invasion status (categorical: low, mid, high), and biomass (continuous, log_10_-transformed) as predictors in separate models for each predictor-response variable combination. For details on these analyses, please see **Appendix Section 4**. We used the R lavaan 0.6-9 (Rosseel 2012) package to construct SEMs testing direct and indirect effects of earthworm invasion on aboveground-arthropod abundance, biomass, and richness, separately (see **Appendix Section 6)**.

## Results

We collected 13,037 aboveground invertebrates (230 Pulmonata individuals included; for brevity, hereafter: arthropods), 4,814 of which were adults. For taxonomic and trophic details, see **Appendix Figures S5 and S6, Table S1**. Arthropod communities differed between invasion-status categories (**Fig. 1**) and along the earthworm-biomass gradient (**Appendix Figure S7**). Out of 18 models testing arthropod responses to earthworm-invasion status, eleven found significant negative relationships, while two relationships were positive (**Fig 1, Table 1**). All three total-arthropod properties responded negatively to earthworm invasion (at least from “low” to “high” invasion). Predator abundance and richness increased with earthworm-invasion status (“mid” to “high”). Out of 18 models testing arthropod responses to increasing earthworm biomass, there were seven significant negative relationships and one significant positive relationship (**Appendix Figure S7, Table S2**). Notably, total-arthropod abundance declined, as well as herbivore abundance and biomass, omnivore abundance, and detritivore abundance, biomass, and richness; only predator biomass increased significantly.

**Table 1:** Results of models relating aboveground-arthropod abundance, biomass, and (morpho)species richness to invasion status (Fig. 1). For each model, the table shows the response variable, arthropod group, sample size (n), model type, response transformation, and p-values for Tukey post hoc and general linear hypotheses tests (see Methods). p-values significant to an alpha level of 0.05 are set in bold. Values are rounded.

| Response | Group | n | Model type | Resp. transf. | p low-high | p low-mid | p mid-high |
| --- | --- | --- | --- | --- | --- | --- | --- |
| Abundance | all | 60 | aov | log10 | < **0.001** | < **0.001** | 0.184 |
| Abundance | herbivores | 60 | aov | log10 | < **0.001** | < **0.001** | 0.137 |
| Abundance | omnivores | 60 | aov | log10 | < **0.001** | **0.040** | **0.030** |
| Abundance | predators | 60 | aov | log10 | 0.682 | 0.238 | **0.043** |
| Abundance | detritivores | 60 | aov | log10 | < **0.001** | < **0.001** | 0.424 |
| Abundance | parasitoids | 60 | aov | log10(+1) | 0.405 | 0.991 | 0.480 |
| Biomass | all | 60 | aov | log10 | **0.042** | 0.800 | 0.166 |
| Biomass | herbivores | 60 | aov | log10 | **0.002** | 0.295 | 0.113 |
| Biomass | omnivores | 60 | aov | log10 | 0.060 | 0.845 | **0.015** |
| Biomass | predators | 60 | aov | log10 | 0.135 | 0.988 | 0.179 |
| Biomass | detritivores | 60 | aov | log10 | < **0.001** | **0.002** | 0.894 |
| Biomass | parasitoids | 60 | aov | log10(+1) | 0.859 | 0.981 | 0.758 |
| Richness | all | 60 | glm.nb | none | **0.025** | 0.058 | 0.942 |
| Richness | herbivores | 60 | glm | none | 0.405 | 0.998 | 0.438 |
| Richness | omnivores | 60 | glm | none | 0.074 | 0.199 | 0.884 |
| Richness | predators | 60 | glm | none | 0.067 | 0.675 | **0.007** |
| Richness | detritivores | 60 | glm | none | < **0.001** | < **0.001** | 0.963 |
| Richness | parasitoids | 60 | glm | none | **0.033** | 0.519 | 0.329 |

**Fig. 1.**
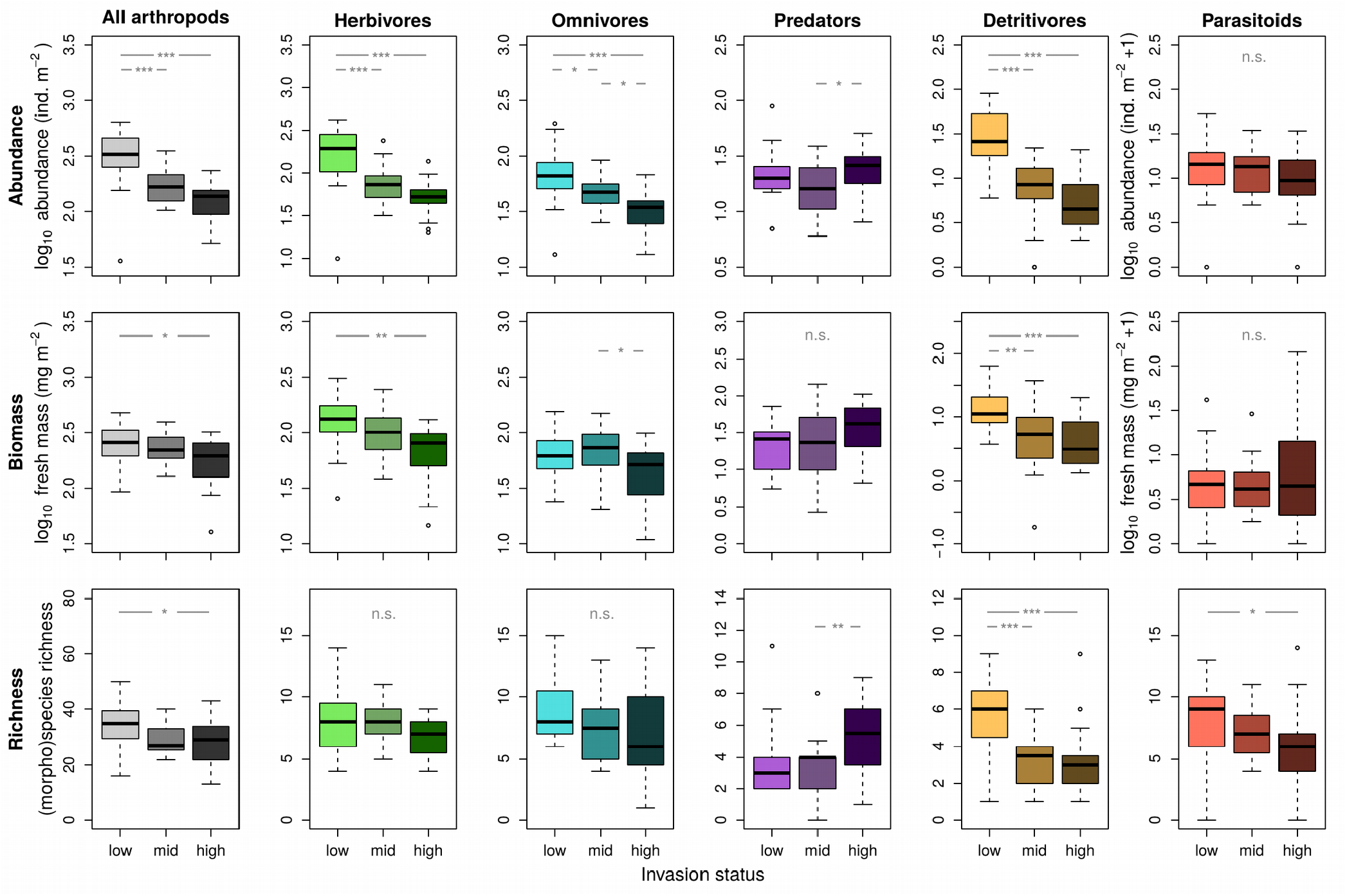
Effects of earthworm invasion status (low, mid, high - lighter to darker color shades) on the abundance (upper row), biomass (middle row) and (morpho)species richness (lower row) of total aboveground arthropods (gray), herbivores (green), omnivores (turquoise), predators (purple), detritivores (brown), and parasitoids (red). Asterisks and “n.s.” illustrate significance levels for differences between invasion-status categories (“n.s.” not significant, p>0.05; ^*^ p<=0.05; ^**^ p<=0.01; ^***^ p<=0.001). P-values are from simple linear models and glm’s with Poisson-distributed response variables (richness models), respectively. N=60. For model results, please see Table 1.

The three SEM’s showed direct and indirect effects of invasive earthworms on aboveground-arthropod communities (**Fig. 2, Tables S3-5**). Earthworm biomass directly increased predator and parasitoid abundance and directly decreased detritivore, herbivore, and omnivore abundance (**Fig. 2b**). It indirectly increased predator abundance via herbivore abundance and indirectly decreased predator and parasitoid abundance via detritivore abundance. Earthworm biomass directly increased predator biomass and directly decreased detritivore and herbivore biomass (**Fig. 2c**). It indirectly decreased predator biomass via detritivore biomass and parasitoid biomass via herbivore biomass. Earthworm biomass directly increased predator richness and directly decreased detritivore richness (**Fig. 2d**). It indirectly decreased predator and parasitoid richness via detritivore richness. There were no significant effects of earthworm biomass on plant cover or richness.

**Fig. 2.**
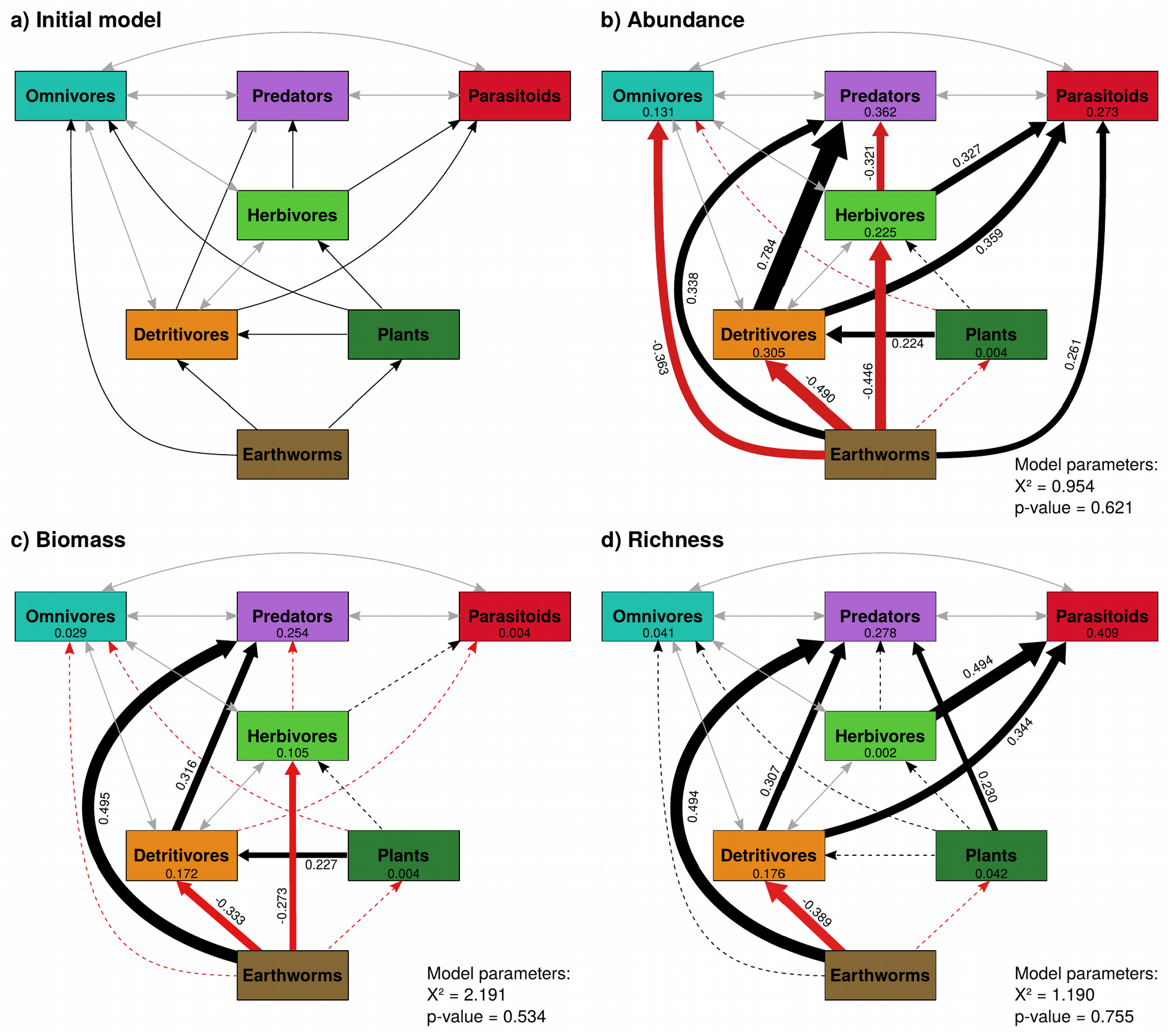
Structural equation models illustrating direct and indirect effects of earthworm invasion on plants and aboveground-arthropod communities. a) Represents the initial model. Final models b-d) were obtained following the steps outlined in **Appendix Section 6**. Brown boxes represent earthworm biomass. Dark green boxes represent plant total cover (b and c), or plant species richness (d). All other boxes represent trophic group abundance (b), biomass (c), or species richness (d). Gray, double-headed arrows show covariances. Black, red, and dashed lines show positive, negative, and non-significant paths, respectively. Numbers next to significant paths are standardized path coefficients. Numbers inside boxes show R^2^ values. N=60.For detailed model outputs, see **Tables S3-S5**.

## Discussion

Our study highlights belowground invasions as a relevant, yet underrepresented driver of aboveground-arthropod decline with impacts cascading across trophic levels. All feeding types and community properties showed significant responses, with only predator communities directly profiting from earthworm invasion in simple models. Our SEMs illustrate how these net positive effects can be decomposed into direct and indirect effects across trophic levels.

In contrast to our expectations but in line with some previous work (e.g. Bohlen et al. 2004, Craven et al. 2017), earthworms had non-significant negative effects on the plant community. The lack of significance might be caused by earthworms changing plant functional diversity and composition instead of total cover and richness (Fleri et al. 2021, Thouvenot et al. 2021) or by high variability in plant communities. Plant cover and species richness supported higher detritivore abundance and biomass, as well as predator richness – supposedly by providing more resources and increased habitat heterogeneity (MacArthur 1972, Gonzalez-Megias et al. 2007). Ubiquitous negative effects of earthworm biomass on detritivores, and omnivore abundance, were likely caused by exploitation competition for litter as a resource strongly diminished by earthworm invasion (Bohlen et al. 2004, Nuzzo et al. 2009) and in this forest particularly (Eisenhauer et al. 2007). Negative effects of earthworm biomass on herbivores might, for example, be caused by earthworm-induced changes in plant secondary metabolites (Thakur et al. 2021), or alternatively via impacts on herbivore life stages in the soil (Ferlian et al. 2018, Jochum et al. 2021).

Across community properties, there were consistent and strong, direct positive effects of earthworm biomass on predators, and on parasitoid abundance, that were not mediated by plant richness or cover, or intermediate trophic levels. Such effects might be mediated by altered habitat structure, such as reduced litter layers (Eisenhauer et al. 2007), or plant-community properties (Fleri et al. 2021), but we need further analyses to better understand the underlying mechanisms. It is likely that these seemingly direct effects are mediated by parameters not included in our models. Detritivores facilitated predators and parasitoids, the former as prey, the latter potentially as a host species, or indirectly via cascading positive effects on plants and herbivores (which we did not test; Megías and Müller (2010)). Herbivores facilitated parasitoids, most prominently in the richness SEM. As herbivore richness was not driven by plant richness, it might respond to plant-functional diversity (Siemann et al. 1998), which could also mediate the direct positive effect of earthworms on parasitoids. Finally, the negative relationship between herbivore and predator abundance might indicate that predators have reduced herbivores (top-down effect) instead of herbivores increasing predators (bottom-up effect; Letourneau et al. 2009).

As one of the first studies reporting effects of invasive earthworms on aboveground-arthropod communities, our paper highlights several topics for future research. First, we need studies investigating the effects of earthworm invasion on vegetation structure, functional diversity, and plant metabolites, as well as their multifaceted impact on arthropod communities (Schuldt et al. 2019, Thakur et al. 2021, Thouvenot et al. 2021). Furthermore, we need to assess the consequences of belowground invasions and the subsequent aboveground arthropod community changes for consumers of arthropods (Lister and Garcia 2018), above-belowground energy flux, ecosystem functions, and services (Barnes et al. 2020, Eisenhauer and Hines 2021, Jochum and Eisenhauer 2021). Future studies should also investigate if earthworm invasion facilitates secondary invasions in aboveground-arthropod communities, potentially facilitated by non-native plants (Craven et al. 2017), and how earthworm invasion might interact with other global-change drivers (Fisichelli et al. 2013, Cameron et al. 2015). Finally, given the varying responses of abundance, biomass, and richness, our results suggest that including multiple community parameters is key when comprehensively assessing the mechanisms of arthropod-community declines under global change.

## Acknowledgements

Svenja Haenzel: coordination. Lotte Horn, Michelle Ives, Morgan Blieske, and Sophia Findeisen: field- and lab-work and data management. Barrier Lake Field Station and Adrienne Cunnings (University of Calgary): accommodation and support. Ian Macdonald: help with identification of local plant species.

## Author contributions

MJ, LT, OF, and NE conceived and designed the study. MJ, LT, OF, RZ, and UP did the field work. BK identified arthropod fauna and assessed their trophic identity. MJ analyzed the data and drafted the manuscript. All authors discussed the results and contributed to writing the manuscript.

## Permits

We thank the Government of Alberta, Canada, for granting access and permits (Alberta Environment and Parks, permit no. 19-260) to do research in the forest at Barrier Lake.

## Code and data accessibility

R-code, data, and a README file are being made available upon submission and will be made publicly-available upon acceptance of the paper.

## Supplementary Information

### 1. Study design, earthworm, and plant communities

**Figure S1.**
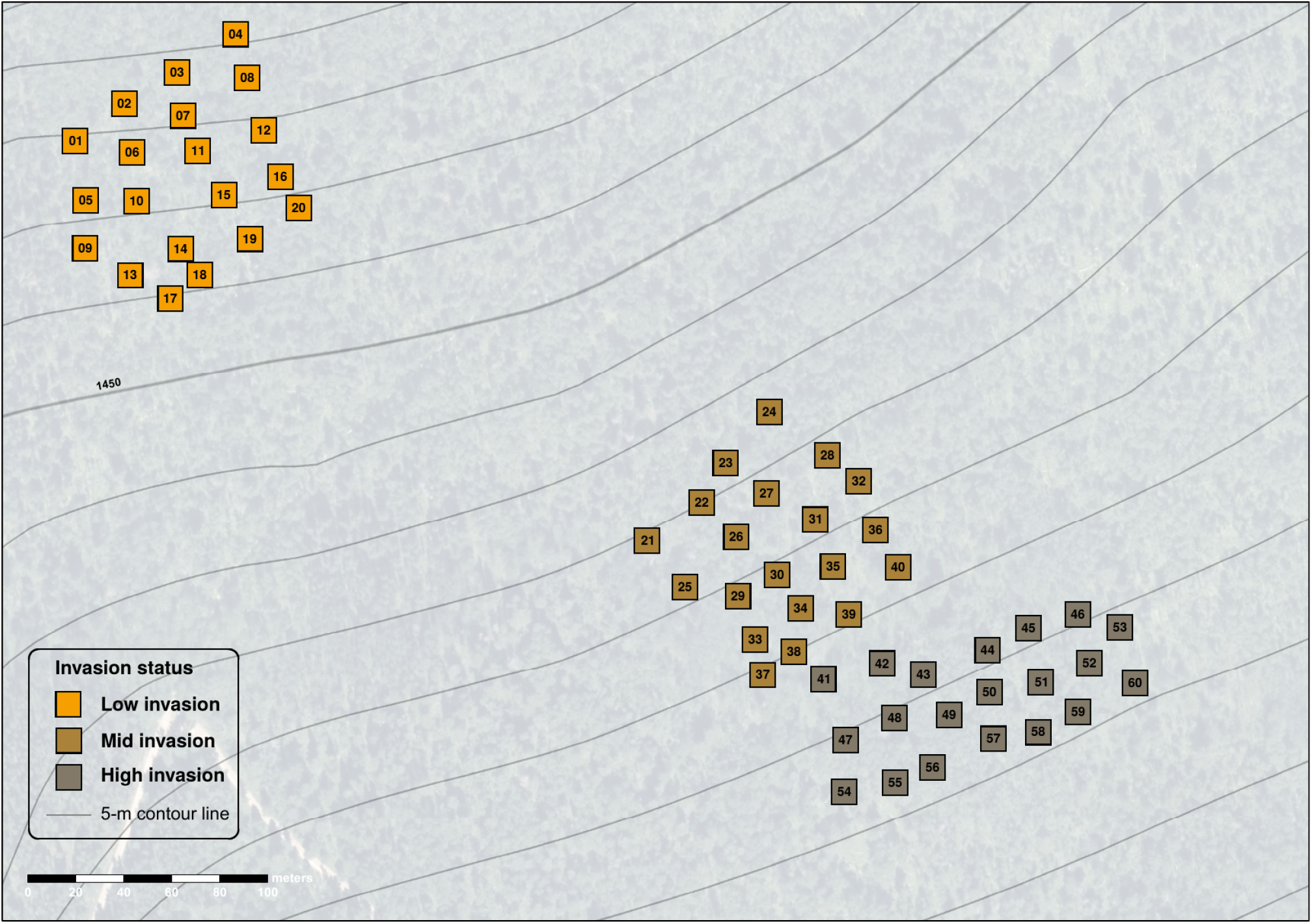
Distribution of 60 study plots on the forest slope above Barrier Lake, Kananaskis Valley, Alberta, Canada. Colors show low (orange), mid (light brown), and high (dark brown) invasion-status.

#### Earthworm communities

Earthworms were extracted on 0.25 m^2^ per plot using a combination of digging and hand-sorting (upper 10 cm), and mustard extraction as described in Jochum et al. (2021). Subsequently, the obtained earthworms were stored in 70% ethanol, taken to the lab, counted, weighed for fresh weight, identified to species, and assigned to one of three ecological groups (epigeic, endogeidc, anecic).

We collected 761 earthworms of five species, across all three ecological groups (anecic, endogeic, epigeic; all European earthworm species). Across earthworm-community properties (i.e., abundance, biomass, richness, and ecological group richness), we found significant differences between the three invasion-status categories, all increasing from low, over medium, to high invasion status (**Figure S2**).

**Figure S2.**
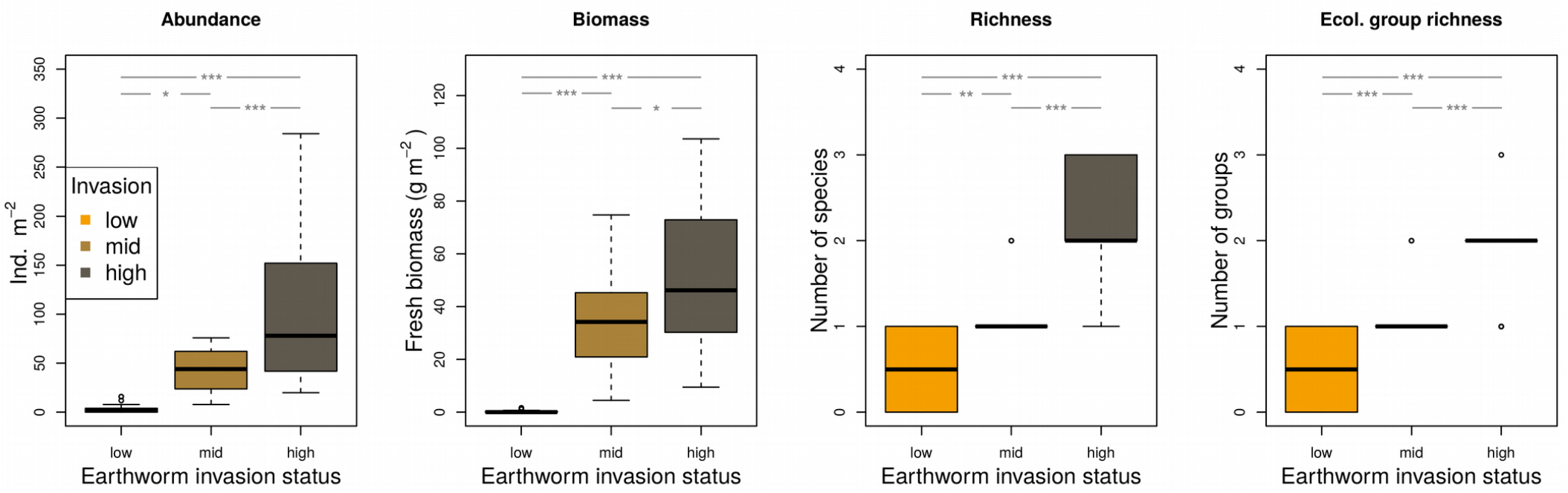
Earthworm community abundance, biomass, richness, and ecological group richness for the three invasion-status categories. Asterisks show significant differences between categories tested with R base function TukeyHSD() on aov() models of community properties against invasion status.

**Figure S3.**
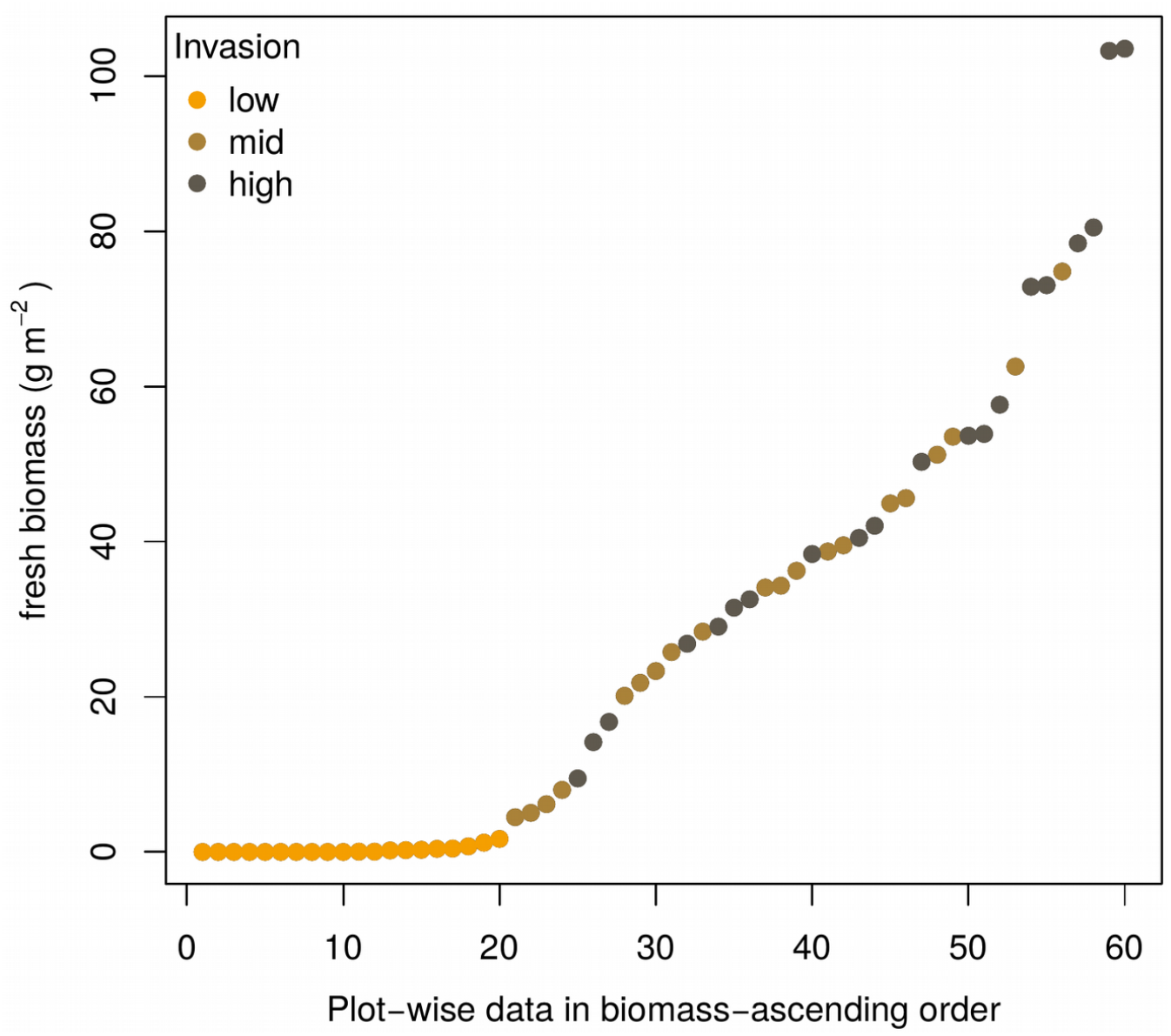
Earthworm biomass for all 60 study plots in increasing order. As can be assumed from the biomass results in Fig. S2, the mid and high invasion status plots are closer to each other and mixed than any of these two groups is with the low status plots.

**Figure S4.**
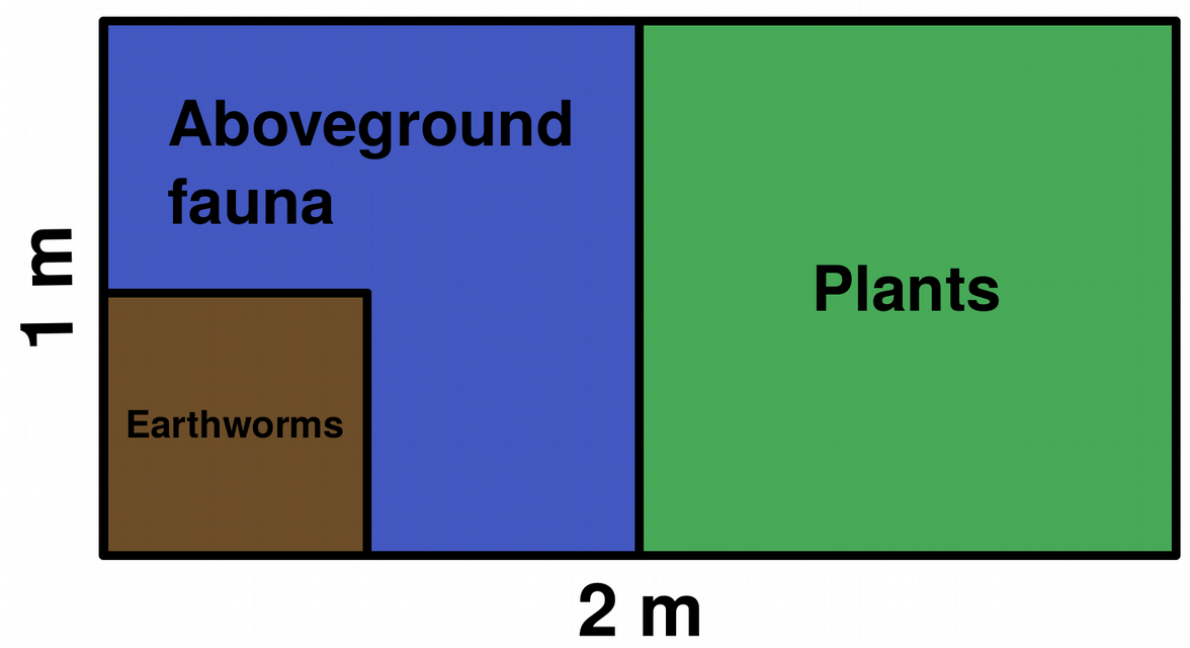
Plot design of the 60 1×2 m plots. We used 1 m^2^ for aboveground fauna sampling and afterwards a quarter m^2^ of the same area for earthworm sampling, and the other 1 m^2^ for plant community assessments.

#### Plant communities

Plant cover was estimated using 13 cover categories (<1%, 1-3%, 3-5%, 5-15%, eight subsequent 10% steps, and >95%). The median values of these categories were used in all analyses. Missing total plant cover for one plot was inferred by averaging over the 19 other plots in this invasion status (high).

### 2. Aboveground-arthropod communities

We sampled aboveground arthropods using an ecoTech (Bonn, Germany) ecoVac insect suction sampler and a 2×1×1 m (lxbxh) insect rearing cage (BugDorm-6E1020, 150×150 µm Nylon mesh, bottom mesh removed) between June 5^th^ and 10^th^ 2019 (covering all three categories on each day). We placed the mesh cage on the ground when approaching the plot. Then, we collected vegetation- and ground-dwelling arthropods by operating the suction sampler across all vegetation and ground with full throttle for 2 min (90° turning the cage after 1 min to better reach all areas) and afterwards emptying out the cage. Suction samples were kept cool and dark in the field, put in a -20 °C freezer overnight, hand-sorted to separate animals from debris, and stored in 70% ethanol. After removing springtails and mites, all remaining animals were then identified to (morpho)species, assigned to a trophic feeding guild. Communities were numerically dominated by Hemiptera, Diptera, Araneae, Thysanoptera, and Hymenoptera, with pure herbivores dominating, followed by predatory-herbivorous arthropods, pure predators, pure detritivores, and parasitoids (**Figures S5 and S6, Table S1)**. Subsequently, up to five individuals per morphospecies-plot combination were measured for body length. Afterwards, we used length-mass regressions to estimate fresh body masses (**Appendix section 3**; Mercer et al. (2001), Wardhaugh (2013), Sohlström et al. (2018)).

**Figure S5.**
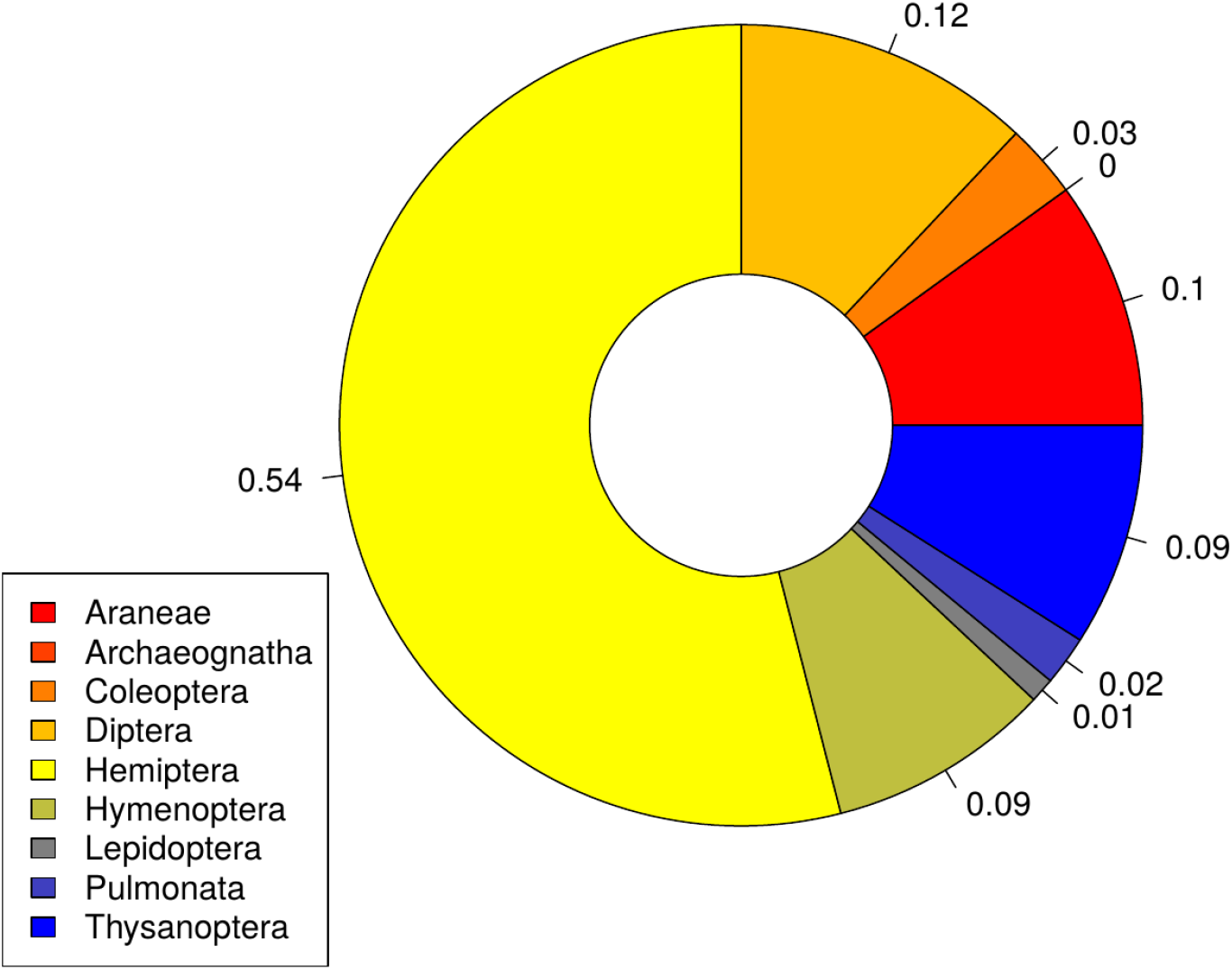
Proportional distribution of taxa among the 13,037 individuals (only groups with >20 ind. shown).

**Figure S6.**
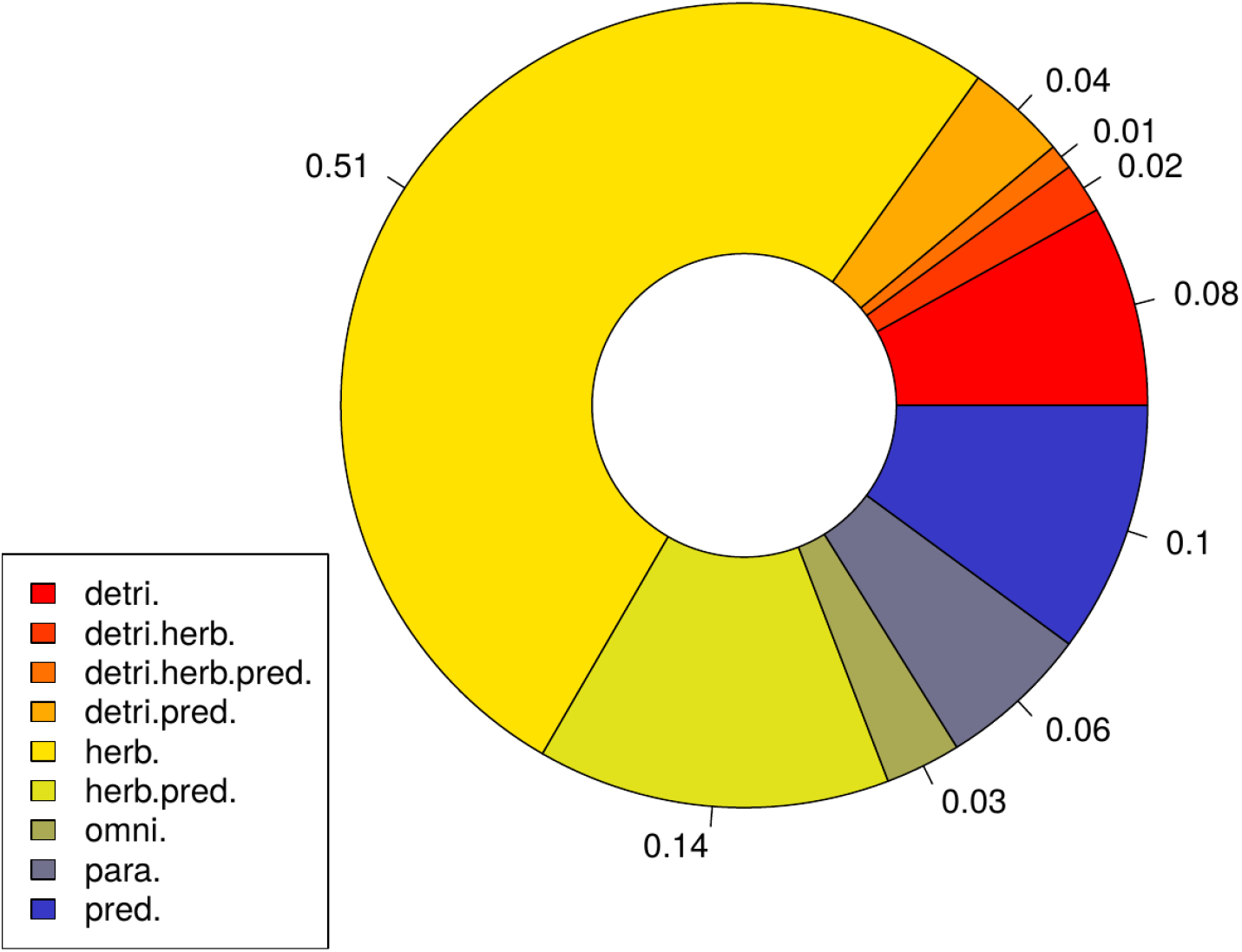
Proportional distribution of trophic feeding guilds among the 13,037 individuals (only groups with >20 ind. shown).

**Table S1:**
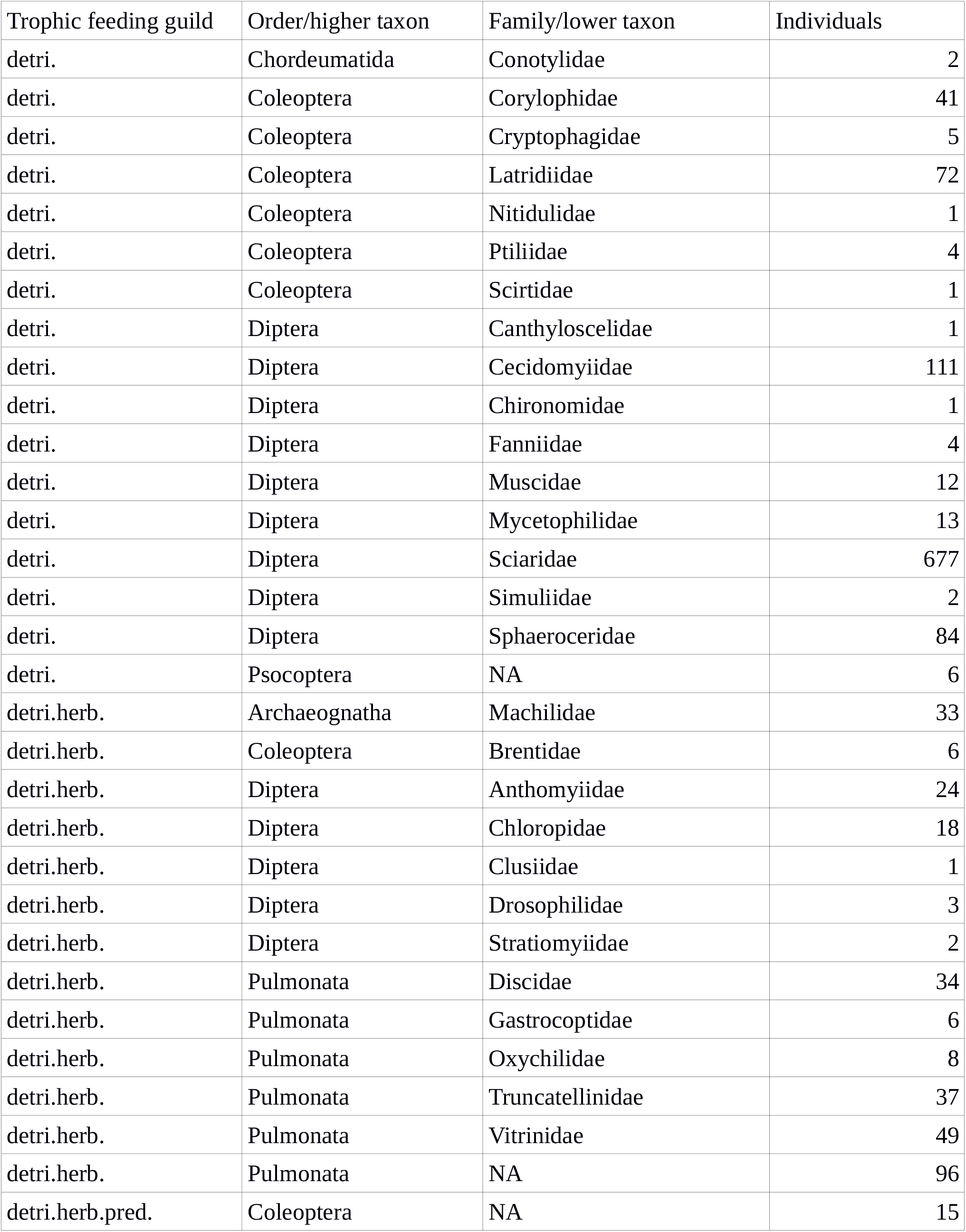
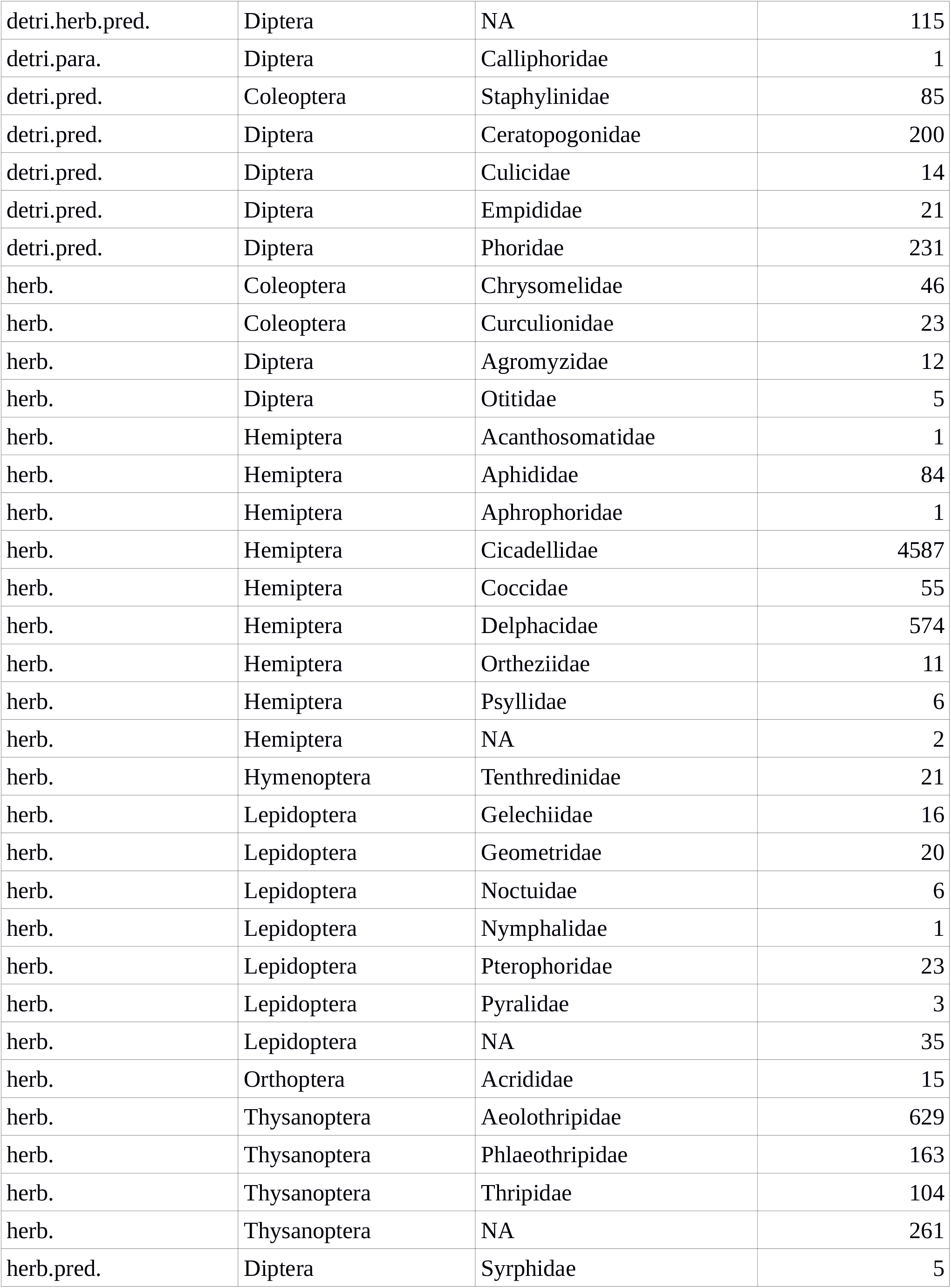
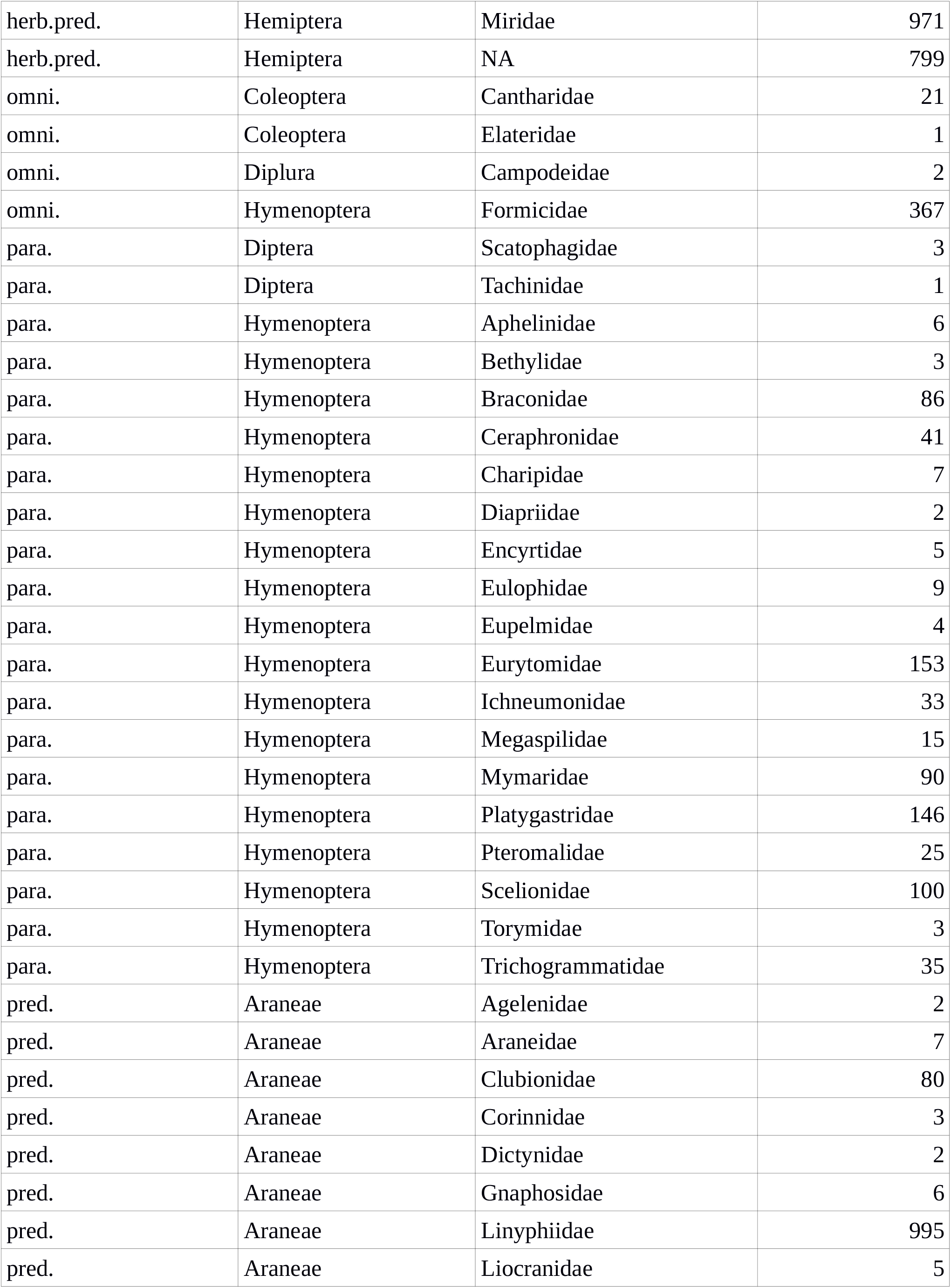
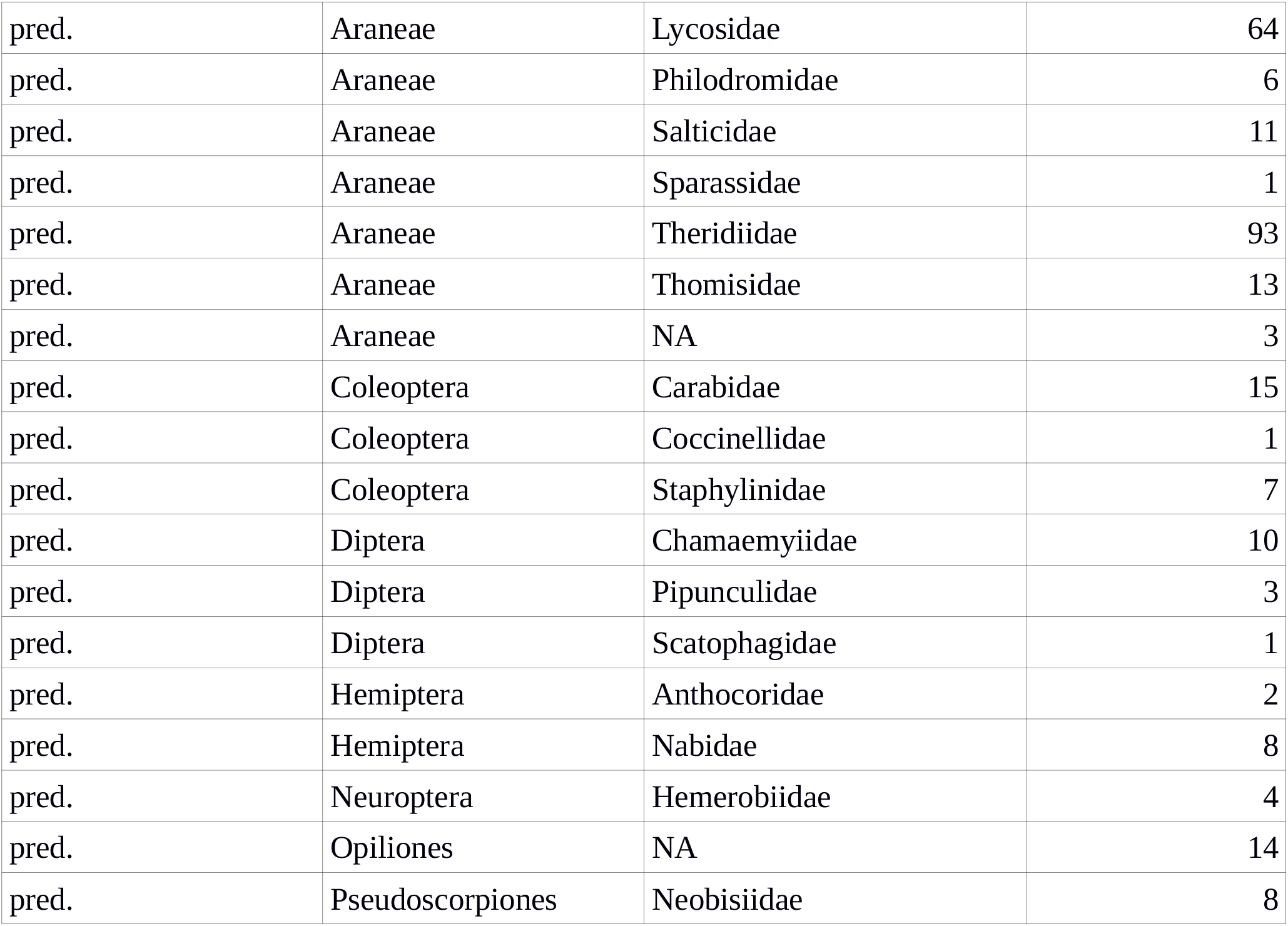
Distribution of the 13,037 individuals across taxonomic family and order (or lower and higher taxonomic levels) for each of the ten trophic feeding guilds. “NA” means that a taxon level was unknown.

### 3. Length-mass regressions

We used body length-fresh body mass regressions from Sohlström et al. (2018) for all taxa except for Pulmonata as follows: LTR model, temperate (Araneae, Chordeumatidae-Polydesmida, Coleoptera, Diptera adults, Hemiptera, Hymenoptera, Lepidoptera adults, Neuroptera, Opiliones, Orthoptera) and tropical (Diptera larvae, Lepidoptera larvae, Pseudoscorpiones, Psocoptera), LR model, temperate (Archaeognatha, Diplura, Thysanoptera). Models with code including “L”, “T”, and “R” use body length (L), taxonomic information (T), and geographical region (R, temperate or tropical). For taxa where specific taxonomic regressions were not available, we used the LR models. For those without temperate models, we used tropical, but taxonomically-specific regressions. Body masses of Pulmonata were calculated using a combination of a length-dry mass regression for gastropods (Wardhaugh 2013) and a general dry-fresh mass conversion (Mercer et al. 2001). For 63 out of 13,037 individuals, there was no length measurement available. These lengths were replaced either a) by averaging over the other individuals measured in the same plot or b), if no other lengths were available in the same plot, by averaging over all individuals of that species from all plots.

### 4. Statistical analysis

We used simple linear models with lm() for earthworm biomass and anova’s with aov() for invasion-status models. For richness models, we used generalized linear models with glm() with “family=poisson” and, if overdispersed (tested using the performance package; Lüdecke et al. 2020; only the case for total richness), we used negative binomial generalized linear models with glm.nb() (MASS package; Venables and Ripley 2002)). We log_10_-transformed abundances and biomasses (log_10_(x+1) for parasitoids because of zero-abundance cases). We used Tukey post hoc tests with TukeyHSD() in base R for lm() and aov() models, and general linear hypotheses tests with glht() in multcomp (Hothorn et al. 2008) for richness glm’s to detect differences between invasion-status categories.

### 5. Models of arthropod community properties versus earthworm biomass

**Fig. S7:**
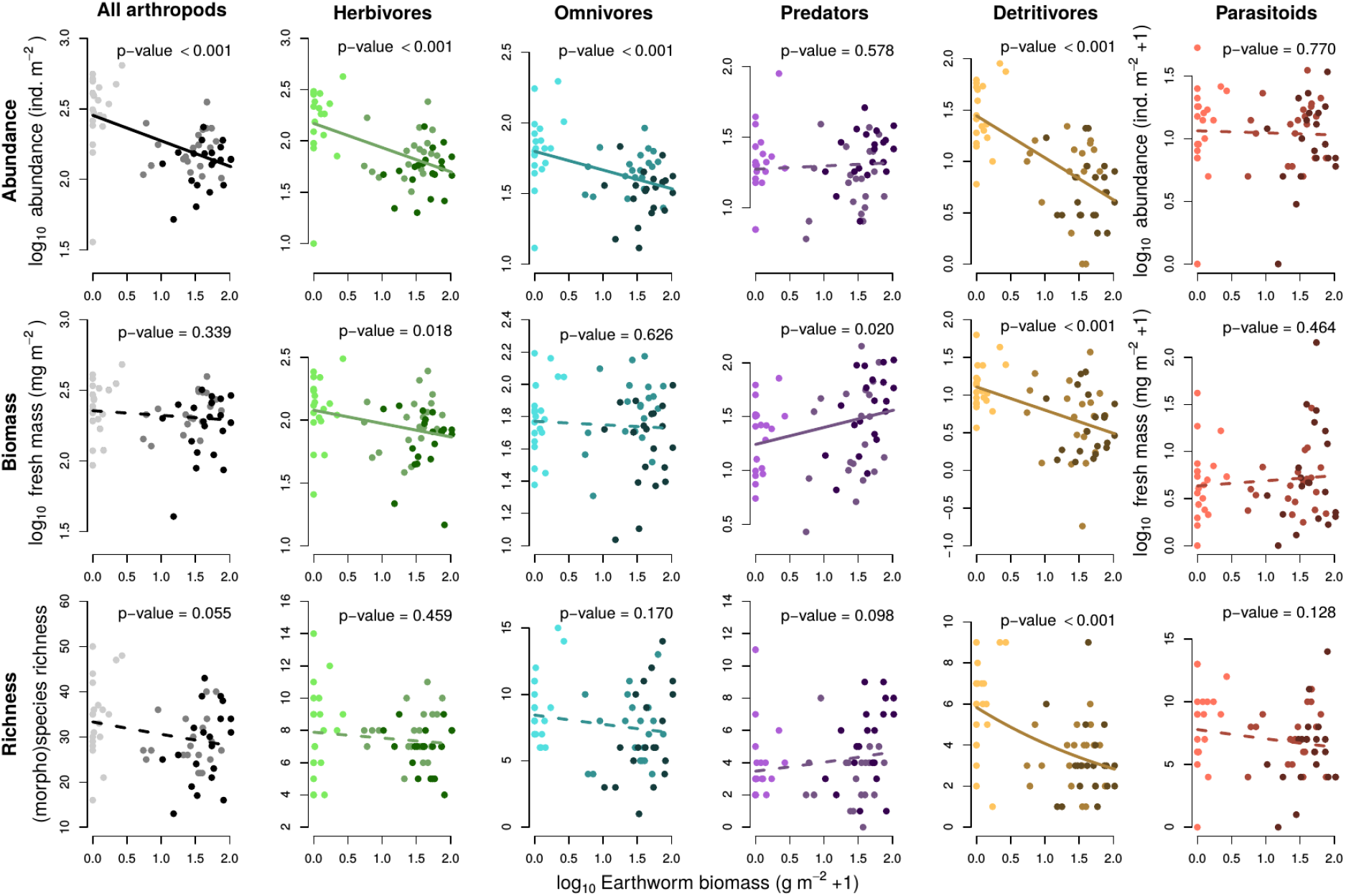
Effects of invasive earthworm biomass on the abundance (upper row), biomass (middle row) and (morpho)species richness (lower row) of aboveground arthropods (all, gray), herbivores (green), omnivores (turquoise), predators (purple), detritivores (brown), and parasitoids (red). Solid and dashed lines show significant and non-significant relationships, respectively. P-values are from simple linear models and glm’s with Poisson-distributed response variables (richness models), respectively. Model predictions in richness models are curved because of the model type. Earthworm biomass, all abundances, and all biomasses have been log_10_ transformed to meet model assumptions. Point colors show low, mid, and high invasion from light to darker shades for each color family. N=60. For model results, please see Table S2.

**Table S2:**
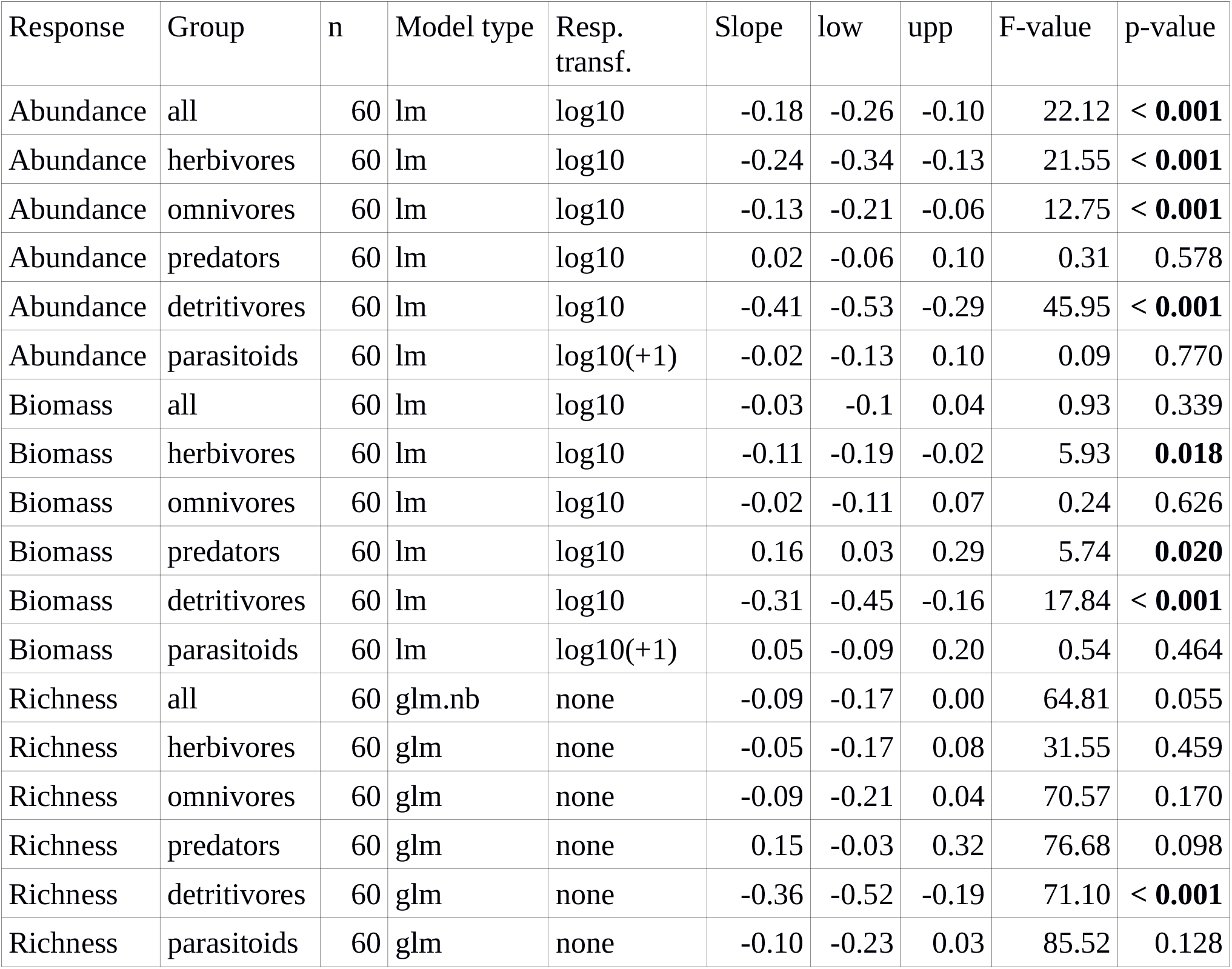
Results of models relating aboveground arthropod abundance, biomass, and (morpho)species richness to earthworm biomass (Fig. S7). For each model, the table shows the response variable, arthropod group, sample size (n), model type, response transformation, slope estimates, approximated lower and upper 95% confidence intervals, F- and p-value. P-values are from simple linear models and glm’s with Poisson distributed response variables (richness models), respectively. Predictor variable is earthworm biomass (log_10_-transformed) for all models. p-values significant to an alpha level of 0.05 are set in bold. Values are rounded.

| Response | Group | n | Model type | Resp. transf. | Slope | low | upp | F-value | p-value |
| --- | --- | --- | --- | --- | --- | --- | --- | --- | --- |
| Abundance | all | 60 | lm | log10 | -0.18 | -0.26 | -0.10 | 22.12 | < **0.001** |
| Abundance | herbivores | 60 | lm | log10 | -0.24 | -0.34 | -0.13 | 21.55 | < **0.001** |
| Abundance | omnivores | 60 | lm | log10 | -0.13 | -0.21 | -0.06 | 12.75 | < **0.001** |
| Abundance | predators | 60 | lm | log10 | 0.02 | -0.06 | 0.10 | 0.31 | 0.578 |
| Abundance | detritivores | 60 | lm | log10 | -0.41 | -0.53 | -0.29 | 45.95 | < **0.001** |
| Abundance | parasitoids | 60 | lm | log10(+1) | -0.02 | -0.13 | 0.10 | 0.09 | 0.770 |
| Biomass | all | 60 | lm | log10 | -0.03 | -0.1 | 0.04 | 0.93 | 0.339 |
| Biomass | herbivores | 60 | lm | log10 | -0.11 | -0.19 | -0.02 | 5.93 | **0.018** |
| Biomass | omnivores | 60 | lm | log10 | -0.02 | -0.11 | 0.07 | 0.24 | 0.626 |
| Biomass | predators | 60 | lm | log10 | 0.16 | 0.03 | 0.29 | 5.74 | **0.020** |
| Biomass | detritivores | 60 | lm | log10 | -0.31 | -0.45 | -0.16 | 17.84 | < **0.001** |
| Biomass | parasitoids | 60 | lm | log10(+1) | 0.05 | -0.09 | 0.20 | 0.54 | 0.464 |
| Richness | all | 60 | glm.nb | none | -0.09 | -0.17 | 0.00 | 64.81 | 0.055 |
| Richness | herbivores | 60 | glm | none | -0.05 | -0.17 | 0.08 | 31.55 | 0.459 |
| Richness | omnivores | 60 | glm | none | -0.09 | -0.21 | 0.04 | 70.57 | 0.170 |
| Richness | predators | 60 | glm | none | 0.15 | -0.03 | 0.32 | 76.68 | 0.098 |
| Richness | detritivores | 60 | glm | none | -0.36 | -0.52 | -0.19 | 71.10 | < **0.001** |
| Richness | parasitoids | 60 | glm | none | -0.10 | -0.23 | 0.03 | 85.52 | 0.128 |

### 6. SEM approach

All three SEMs used earthworm biomass as the basal exogenous variable. Plants are represented by total cover for abundance and biomass SEMs, and plant species richness for the richness SEM. All other groups are represented by their abundance, biomass, and richness in the respective models.

#### a) Initial model

The initial model (see Fig. 1a) was based on assumed trophic relationships, covariance for groups sharing resources or sharing a joint feeding type (omnivores with most others). Earthworms were additionally allowed to impact plants directly (Scheu 2001, Van Groenigen et al. 2014).

Assumed trophic relationships: Detritivores consume dead plant material. Herbivores consume plants. Omnivores feed on plants. Predators consume detritivores and herbivores. Parasitoids feed on herbivores and detritivores.

Groups sharing resources: Omnivores and earthworms. Detritivores and earthworms. Predators and Parasitoids. Parasitoids and Omnivores. Omnivores and Predators. Herbivores and detritivores.

Groups with partly overlapping feeding type: Omnivores and detritivores, herbivores, predators, parasitoids.

#### b) Adding paths

We used modification indices (parameter mi) as indicated by the lavaan R package to look for paths that should be added and included those showing values larger than 3.84 as suggested by James Grace in his lavaan tutorials (https://www.usgs.gov/centers/wetland-and-aquatic-research-center/science/quantitative-analysis-using-structural-equation?qt-science_center_objects=0#qt-science_center_objects).

#### c) Selection of final model

Once paths were added, we checked the validity of the model (chi-square statistic and value). If paths were borderline, in our case only for the abundance model, adding the path from earthworm biomass to parasitoid abundance, we compared the models without and with this path using R functions anova() and aictab.lavaan() downloaded from http://jarrettbyrnes.info/ubc_sem/lavaan_materials/ looking at the significance of chisquare tests and AICc differences. In this case, the model with the path added was significantly better than the one without it, so we included it in the final abundance SEM (Fig. 2 b).

We did not deliberately strip down models to improve fit by removing non-significant paths from valid models, because the initially-selected paths are ecologically meaningful, e.g. predators depend on herbivores as a resource even if this is not significant in a given model.

#### d) lavaan output tables for final models

**Table S3:** lavaan standardizedSolution() output for abundance SEM, Fig. 2b. Lhs and rhs denote the left-hand and right-hand side of regression (op ∼) or covariance relationships (op ∼∼). est.std are standardized parameter estimates. se is the corresponding standard error. z is the z-value. P-values of significant paths in bold. Finally, ci.lower and ci.upper show lower and upper 95% confidence intervals for standardized parameters.

| lhs | op | rhs | est.std | se | z | p-value | ci.lower | ci.upper |
| --- | --- | --- | --- | --- | --- | --- | --- | --- |
| plant_cover | ~ | worm_biomass | -0.0658 | 0.1284 | -0.5127 | 0.6082 | -0.3175 | 0.1858 |
| detri_abund | ~ | plant_cover | 0.2242 | 0.106 | 2.1157 | **0.0344** | 0.0165 | 0.4318 |
| detri_abund | ~ | worm_biomass | -0.4901 | 0.0891 | -5.5002 | **< 0.0001** | -0.6648 | -0.3155 |
| herb_abund | ~ | worm_biomass | -0.4458 | 0.0971 | -4.5919 | **< 0.0001** | -0.6361 | -0.2555 |
| herb_abund | ~ | plant_cover | 0.1346 | 0.1131 | 1.1897 | 0.2342 | -0.0871 | 0.3563 |
| pred_abund | ~ | worm_biomass | 0.3385 | 0.1163 | 2.9113 | **0.0036** | 0.1106 | 0.5664 |
| pred_abund | ~ | herb_abund | -0.3208 | 0.13 | -2.4679 | **0.0136** | -0.5756 | -0.066 |
| pred_abund | ~ | detri_abund | 0.7841 | 0.1165 | 6.7306 | **< 0.0001** | 0.5558 | 1.0125 |
| omni_abund | ~ | plant_cover | -0.0287 | 0.1107 | -0.2596 | 0.7952 | -0.2456 | 0.1882 |
| omni_abund | ~ | worm_biomass | -0.3626 | 0.1087 | -3.3355 | **0.0009** | -0.5756 | -0.1495 |
| para_abund | ~ | worm_biomass | 0.2607 | 0.1266 | 2.0596 | **0.0394** | 0.0126 | 0.5087 |
| para_abund | ~ | herb_abund | 0.3273 | 0.1367 | 2.3949 | **0.0166** | 0.0594 | 0.5951 |
| para_abund | ~ | detri_abund | 0.3593 | 0.1403 | 2.5605 | **0.0105** | 0.0843 | 0.6344 |
| detri_abund | ~~ | omni_abund | 0.6229 | 0.0786 | 7.9278 | **< 0.0001** | 0.4689 | 0.7769 |
| herb_abund | ~~ | omni_abund | 0.4543 | 0.1024 | 4.4372 | **< 0.0001** | 0.2536 | 0.655 |
| pred_abund | ~~ | omni_abund | 0.3979 | 0.0835 | 4.7661 | **< 0.0001** | 0.2343 | 0.5616 |
| omni_abund | ~~ | para_abund | 0.186 | 0.0947 | 1.9647 | 0.0495 | 0.0004 | 0.3715 |
| pred_abund | ~~ | para_abund | 0.1935 | 0.1243 | 1.5572 | 0.1194 | -0.05 | 0.4371 |
| detri_abund | ~~ | herb_abund | 0.4613 | 0.1016 | 4.5399 | **< 0.0001** | 0.2622 | 0.6605 |

**Table S4:** lavaan standardizedSolution() output for biomass SEM, Fig. 2c. Lhs and rhs denote the left-hand and right-hand side of regression (op ∼) or covariance relationships (op ∼∼). est.std are standardized parameter estimates. se is the corresponding standard error. z is the z-value. P-values of significant paths in bold. Finally, ci.lower and ci.upper show lower and upper 95% confidence intervals for standardized parameters.

| lhs | op | rhs | est.std | se | z | p-value | ci.lower | ci.upper |
| --- | --- | --- | --- | --- | --- | --- | --- | --- |
| plant_cover | ~ | worm_biomass | -0.0658 | 0.1284 | -0.5127 | 0.6082 | -0.3175 | 0.1858 |
| detri_biomass | ~ | plant_cover | 0.2267 | 0.115 | 1.9704 | **0.0488** | 0.0012 | 0.4522 |
| detri_biomass | ~ | worm_biomass | -0.3332 | 0.1082 | -3.0795 | **0.0021** | -0.5452 | -0.1211 |
| herb_biomass | ~ | worm_biomass | -0.2727 | 0.1157 | -2.3576 | **0.0184** | -0.4995 | -0.046 |
| herb_biomass | ~ | plant_cover | 0.1574 | 0.121 | 1.3005 | 0.1934 | -0.0798 | 0.3946 |
| pred_biomass | ~ | worm_biomass | 0.4953 | 0.102 | 4.8556 | < **0.0001** | 0.2954 | 0.6952 |
| pred_biomass | ~ | herb_biomass | -0.126 | 0.1238 | -1.0184 | 0.3085 | -0.3686 | 0.1165 |
| pred_biomass | ~ | detri_biomass | 0.3164 | 0.1238 | 2.5548 | **0.0106** | 0.0737 | 0.5591 |
| omni_biomass | ~ | plant_cover | -0.1551 | 0.1236 | -1.2547 | 0.2096 | -0.3973 | 0.0872 |
| omni_biomass | ~ | worm_biomass | -0.084 | 0.1268 | -0.6623 | 0.5077 | -0.3325 | 0.1645 |
| para_biomass | ~ | herb_biomass | 0.0367 | 0.1417 | 0.259 | 0.7956 | -0.241 | 0.3144 |
| para_biomass | ~ | detri_biomass | -0.0709 | 0.1415 | -0.5013 | 0.6161 | -0.3482 | 0.2063 |
| detri_biomass | ~~ | omni_biomass | 0.1971 | 0.1239 | 1.5901 | 0.1118 | -0.0458 | 0.44 |
| herb_biomass | ~~ | omni_biomass | 0.2995 | 0.1175 | 2.5497 | 0.0108 | 0.0693 | 0.5297 |
| pred_biomass | ~~ | omni_biomass | 0.2027 | 0.1174 | 1.7266 | 0.0842 | -0.0274 | 0.4329 |
| omni_biomass | ~~ | para_biomass | -0.0133 | 0.1224 | -0.1089 | 0.9132 | -0.2533 | 0.2266 |
| pred_biomass | ~~ | para_biomass | -0.0052 | 0.1291 | -0.0404 | 0.9678 | -0.2582 | 0.2478 |
| detri_biomass | ~~ | herb_biomass | 0.3289 | 0.1151 | 2.8566 | 0.0043 | 0.1032 | 0.5546 |

**Table S5:**
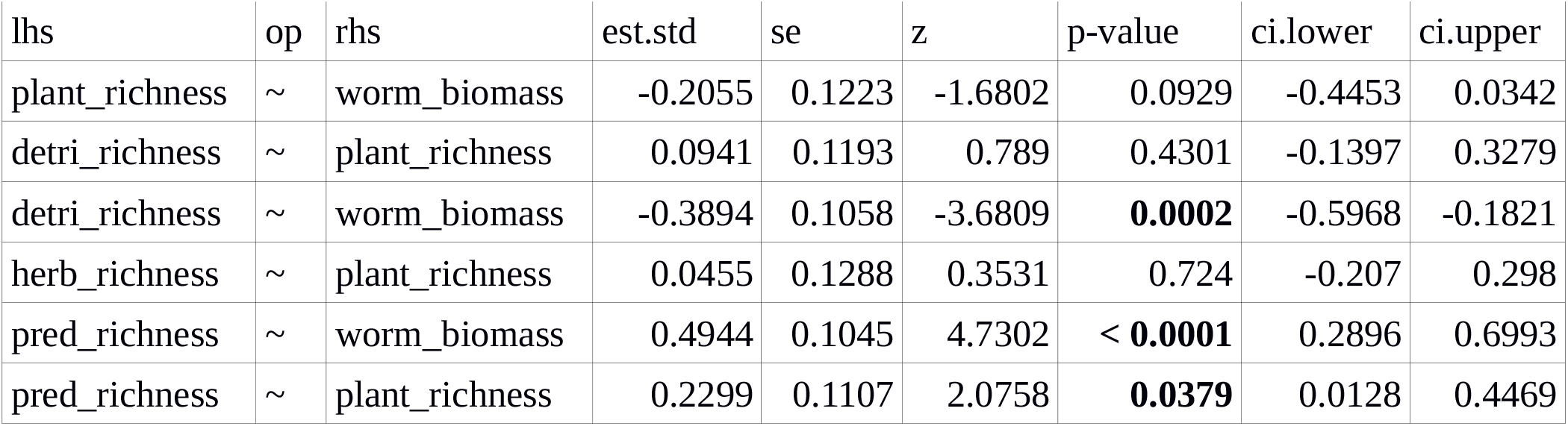
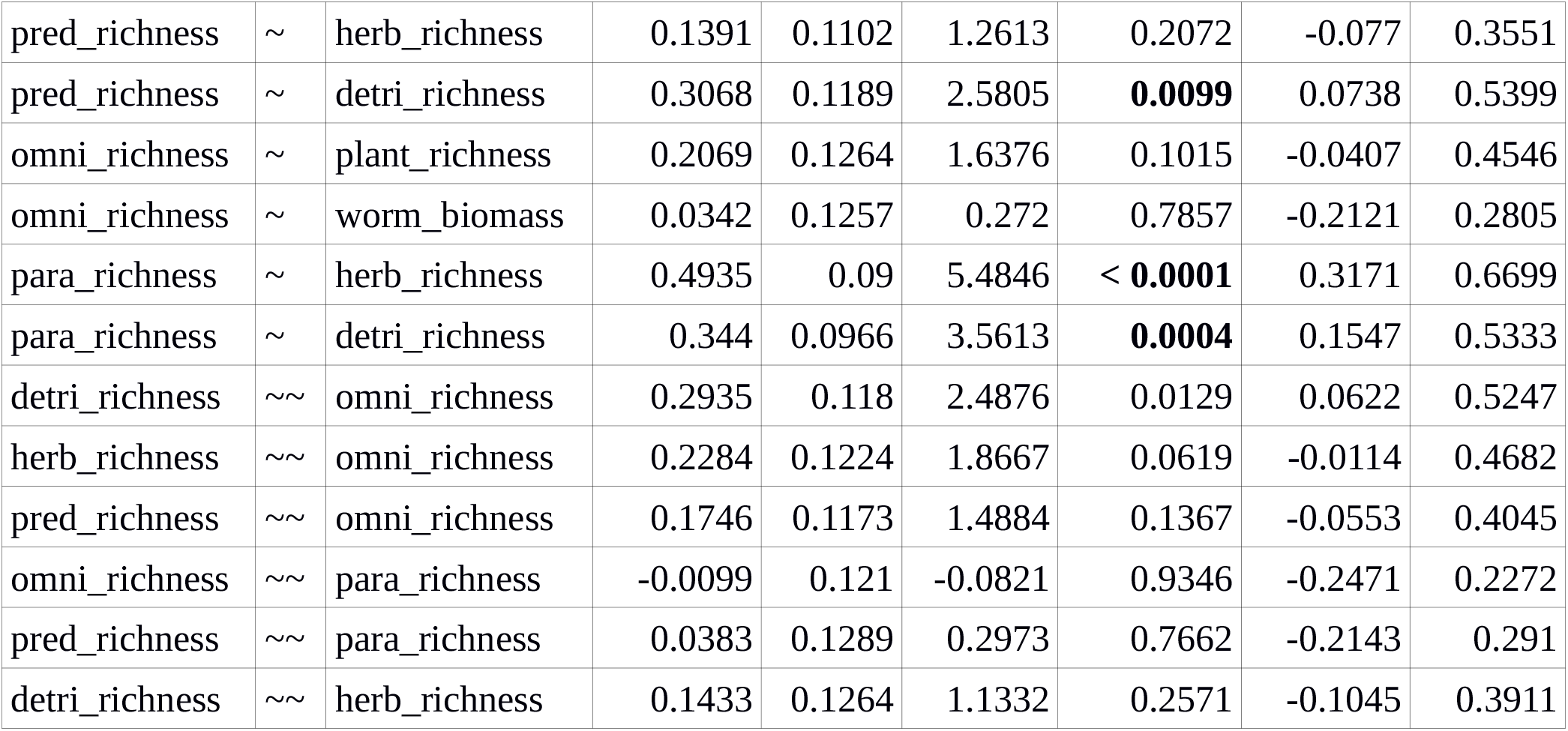
lavaan standardizedSolution() output for richness SEM, Fig. 2d. Lhs and rhs denote the left-hand and right-hand side of regression (op ∼) or covariance relationships (op ∼∼). est.std are standardized parameter estimates. se is the corresponding standard error. z is the z-value. P-values of significant paths in bold. Finally, ci.lower and ci.upper show lower and upper 95% confidence intervals for standardized parameters.

